# The ASNS inhibitor ASX-173 potentiates L-asparaginase anticancer activity

**DOI:** 10.1101/2025.07.03.662851

**Authors:** Victor Tatarskiy, Wai-Kin Chan, Lin Tan, Alvina Khamidullina, Iqbal Mahmud, Shwetha V. Kumar, Julia Nuzhina, Sara A. Martinez, Bao Q. Tran, Anna Skwarska, Nataliya Pavlenko, Alena Dorofeeva, Marina Konopleva, Dmitry Genis, John N. Weinstein, Roman Kombarov, Philip L. Lorenzi

**Author notes:** Equal contribution (Victor Tatarskiy, Wai-Kin Chan).

## Abstract

Cancer cells reprogram metabolic pathways to meet increased energy and biosynthetic demands. Among those pathways, elevated asparagine metabolism regulated by asparagine synthetase (ASNS) has been linked to tumor progression in various cancers, driving cell proliferation, chemoresistance, and metastasis. ASNS inhibition represents a promising therapeutic strategy, but inhibitors have shown limited efficacy due to poor specificity and cell permeability. Through phenotypic screening, we identified ASX-173, a cell-permeable small molecule that inhibits ASNS at nanomolar concentrations. Biochemical and cellular assays confirm the specificity of ASX-173 activity and demonstrate its potentiation of the anti-cancer activity of L-asparaginase (ASNase), a key component of childhood acute lymphoblastic leukemia therapy. Mechanistically, the combination treatment disrupted nucleotide synthesis and induced cell cycle arrest and apoptosis. In a mouse model of acute myeloid leukemia, the combination significantly delayed the growth of OCI-AML2 xenografts. Analysis of data from The Cancer Genome Atlas (TCGA) revealed that *ASNS* mRNA expression is associated with poor survival in some cancer types and that ASNS protein levels are elevated in multiple solid tumors compared with the levels in normal tissues, suggesting possible broad utility of ASNS inhibition across the landscape of cancer. Together, these findings establish ASX-173 as a promising ASNS inhibitor and, for the first time, demonstrate a viable strategy to target ASNS therapeutically—an approach that has long remained elusive.

## INTRODUCTION

Cancer cells reprogram their metabolism to sustain rapid proliferation, traditionally viewed through the lens of altered glucose metabolism and the Warburg effect^1^. Among other metabolic drivers of cancer progression, amino acid metabolism presents notable metabolic vulnerabilities^2^. For example, glutamine’s role in energy production, redox balance, and biosynthesis is well established and targetable with glutaminase (GLS) inhibitors^3^. Asparagine metabolism has been studied as a therapeutic target for over 50 years, but, like glutamine metabolism, targeting it has produced minimal success in the broad oncology setting^4,5^.

The therapeutic potential of targeting asparagine metabolism was first demonstrated by L-asparaginase (ASNase) treatment of childhood acute lymphoblastic leukemia (ALL), for which it is a component of standard therapy. ALL cells are generally deficient in asparagine synthetase (ASNS), rendering them dependent on extracellular asparagine and making them vulnerable to ASNase^6^. However, most cancers rapidly acquire resistance to ASNase through ASNS upregulation, which enables cells to synthesize asparagine intracellularly to counteract the extracellular depletion of asparagine by ASNase^7^. Although ASNase-based therapy has been successful in treating childhood ALL, similar success has not been achieved in adult ALL or other cancer types. Nonetheless, the expanding understanding of asparagine’s roles in cancer biology, including regulation of cellular signaling networks, function as an amino acid exchange factor^8^, and promotion of epithelial-to-mesenchymal transition (EMT)—a critical step in metastasis^9^, continues to highlight asparagine’s therapeutic relevance. Hence, there is renewed interest in targeting asparagine metabolism across multiple cancer types.

Despite the compelling rationale for targeting ASNS therapeutically, however, attempts to develop ASNS inhibitors have faced significant challenges. First-generation sulfoximine-based inhibitors exhibited limited cell penetration^10^. Subsequent adenylate-based inhibitors improved penetration but suffered from poor stability and specificity^11,12^.

Here, we present ASX-173, a small-molecule ASNS inhibitor that exhibits superior cell permeability, target specificity, and potency. Its combination with ASNase achieves a bicompartmental blockade of asparagine metabolism that results in synergistic anticancer activity against multiple cancer cell types, including those resistant to ASNase alone. Notably, ASX-173 renders ASNase-resistant tumors ASNase-sensitive *in vivo*. Our findings establish ASX-173 as a promising therapeutic candidate that warrants further testing in combination with ASNase in childhood ALL, adult ALL, and a broader range of cancer types, and further investigation is warranted to explore its potential in combination with agents beyond ASNase.

## RESULTS

### Phenotypic screening identifies ASX-173 as a context-dependent inhibitor of cancer cell growth

Through an unbiased phenotypic screen for inhibitors of the Wnt/β-catenin pathway, we identified several small molecules that inhibited TCF4/LEF luciferase reporter activity in Wnt-dependent colon cancer cell lines (SW620 and HCT-116). ASX-173 emerged as the most potent of the compounds, exhibiting the lowest IC50 values across multiple cell lines (**Figure 1A** and **Supplemental Figure S1**). Real-time PCR analysis confirmed that ASX-173 treatment suppressed transcription of several endogenous Wnt target genes, including *AXIN1, DKK1, CD133/PROM1*, and *MYC* in HCT-116 cells (**Figure 1B**).

**Figure 1.**
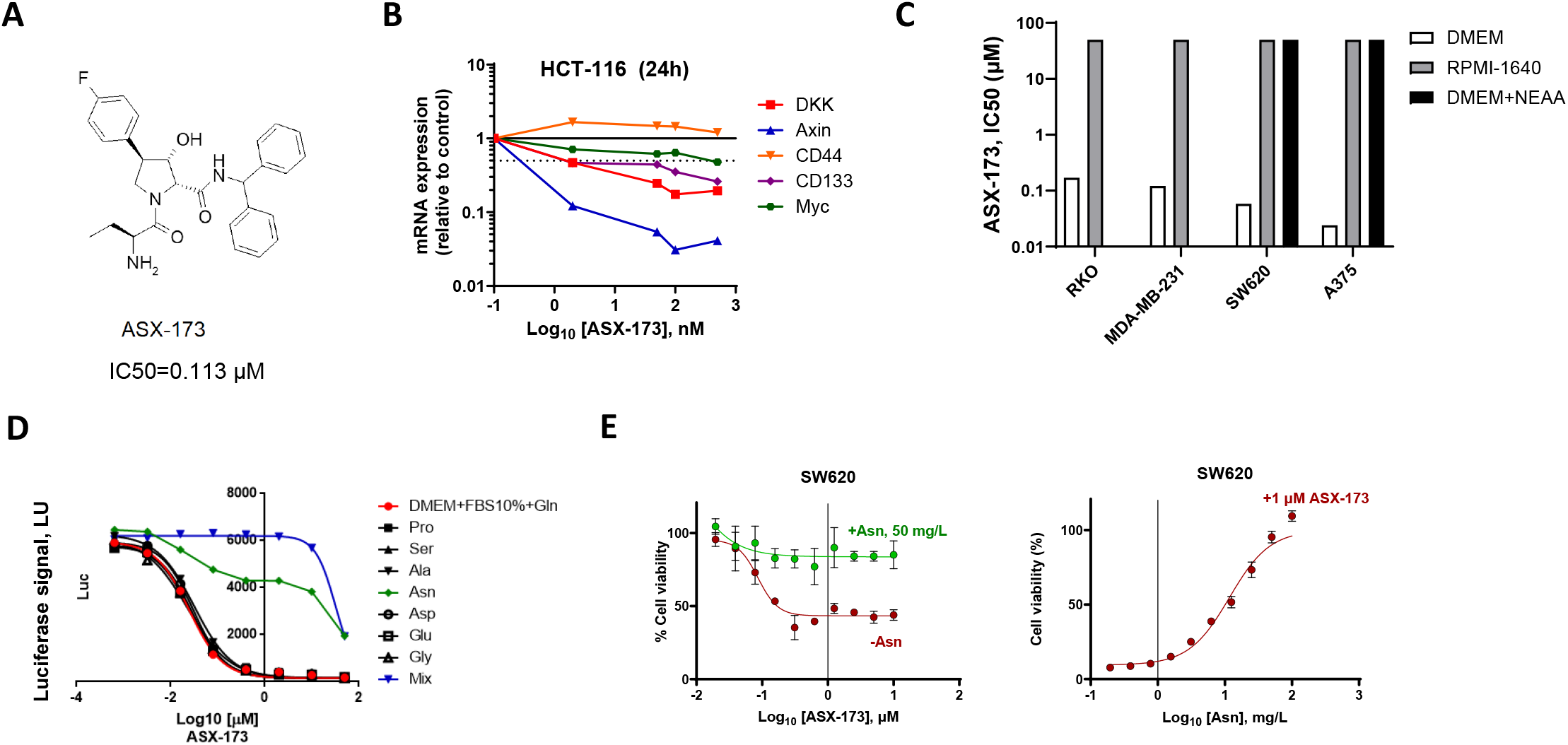
ASX-173 is an inhibitor of cancer cell growth under asparagine-deficient conditions. (**A**) Chemical structure of ASX-173. (**B**) ASX-173 treatment suppressed the expression of endogenous Wnt target genes *AXIN1, DKK1, CD133/PROM1*, and *MYC* relative to control in HCT116 cells. mRNA levels were measured by real-time PCR after treatment with the indicated concentrations of ASX-173 (n=3). *CD44*, a Wnt-independent gene, served as a negative control. Data represent mean fold change relative to the vehicle controls. (**C**) IC_50_ values of ASX-173 in the indicated cell lines cultured in DMEM, RPMI-1640, or DMEM supplemented with NEAA, measured using the MTT assay. (**D**) SW620 cells were treated with ASX-173 at various concentrations in DMEM supplemented with NEAA or specific individual amino acids. Cell viability was measured using the MTT assay. (**E**) SW620 cells were treated with ASX-173 with or without addition of 5 mM asparagine. The MTT assay was performed as in (**D**).

Structure-activity relationship analysis revealed that the initial hits and their derivatives shared a critical substructural feature that correlated with their anticancer activity (**Supplemental Figure S1**). Intriguingly, that activity was strictly dependent on the culture medium composition; compounds exhibited potent cytotoxicity in DMEM medium but minimal to no effect when cells were cultured in RPMI-1640 (**Figure 1C**). A systematic comparison of the composition of the media identified four amino acids as potential mediators of the differential response—asparagine, aspartic acid, glutamic acid, and proline.

To identify the specific amino acid responsible for the effect, we individually supplemented DMEM with each candidate amino acid, including serine, glycine, and alanine as controls. Only asparagine supplementation was capable of abolishing the inhibitory effect of ASX-173 on TCF4/LEF reporter activity (**Figure 1D**), suggesting that ASX-173 specifically interferes with asparagine production. Dose-response studies in SW620 cells demonstrated that supplementation with 11.7 mg/L L-asparagine, corresponding to its IC_50_, partially rescued cell viability, and 50 mg/L L-asparagine provided complete protection against ASX-173-mediated cytotoxicity (**Figure 1E**).

### ASX-173 acts as a potent and selective ASNS inhibitor

To elucidate the molecular target of ASX-173, we leveraged the DepMap database to identify genes whose loss of function mimicked the compound’s amino acid-dependent effects. Analysis of CRISPR and RNAi screens revealed that asparagine synthetase (ASNS) knockout or knockdown exhibited the strongest synthetic lethality under conditions of non-essential amino acid (NEAA) deficiency (**Figure 2A, Supplemental Figures S2A**). Further examination of CRISPR knockout or RNAi knockdown scores demonstrated a robust negative correlation between NEAA deprivation and heightened sensitivity to ASNS deletion (**Figure 2B and Supplemental figures S2B**-**C**), suggesting ASNS as the potential molecular target of ASX-173.

**Figure 2.**
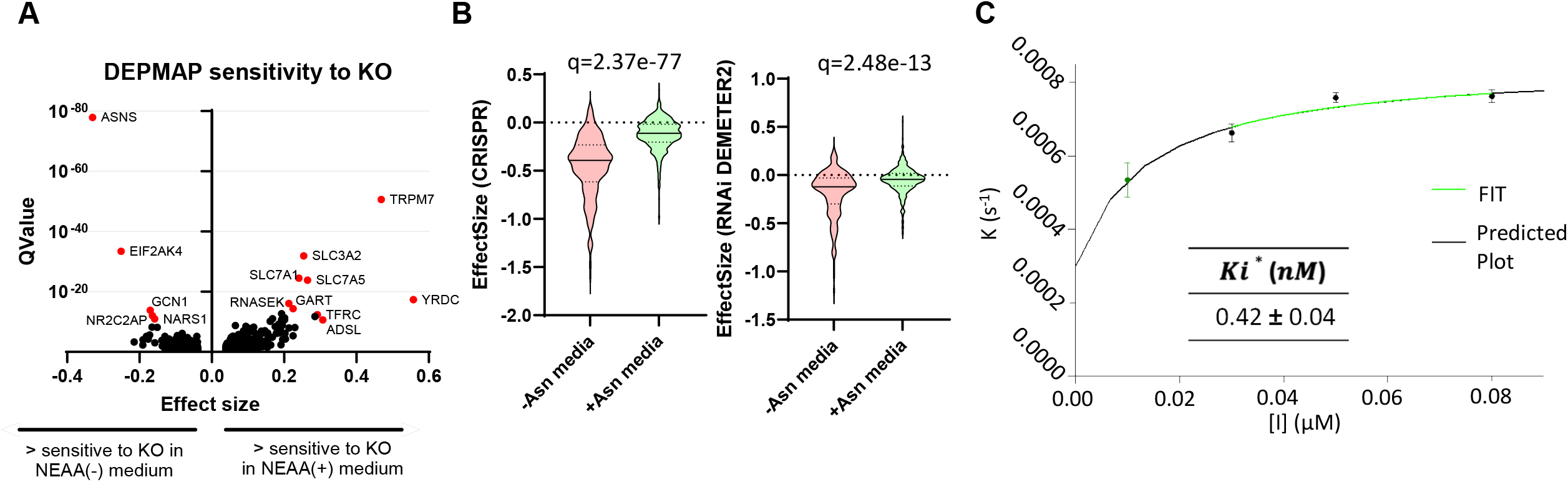
ASX-173 is a potent, selective ASNS inhibitor. (**A**) Volcano plot depicting gene dependency in human cancer cell lines (CRISPR knockout) cultured in NEAA-deficient medium, based on DepMap data. NEAA-containing medium was used as a control. (**B**) Sensitivity violin plot of ASNS-deficient cancer cells (Left panel: CRISPR knockout; Right panel: RNAi knockdown) to NEAA-null (Asn-null) culture medium, as observed in the DepMap database. Asparagine containing medium was used as a control. 35.9% of cell lines in Asn-null media were sensitive to ASNS KO, compared to 3.6% in control media (q=2.37e-77); and 12.3% were sensitive to ASNS KD, compared to 1.3% in control (q=2.48e-13). (**C**) ASNS enzyme kinetics in the presence of ASX-173. Inorganic pyrophosphate (PPI) production was measured in a recombinant human ASNS-mediated asparagine synthesis assay at varying ASX-173 concentrations, and the inhibitory constant (k_i_) was determined as described in Supplementary method II.

To validate that hypothesis, we performed direct binding studies using purified human ASNS protein. We monitored the rate of inorganic pyrophosphate (PPi) production, a byproduct of the ASNS-catalyzed reaction, in the presence of varying concentrations of ASX-173. These biochemical studies revealed ASX-173 as a highly potent ASNS inhibitor with an inhibition constant (k_i_) of 0.4 nM (**Figure 2C**).

### ASX-173 potentiates L-asparaginase activity across diverse cancer types

To validate the specificity of ASX-173 for ASNS, we used a paired cell line model consisting of ASNS-deficient RS4;11 cells (RS4;11) and their ASNS-expressing counterpart (RS4;11_ASNS), which were derived from RS4;11 cells cultured in the absence of asparagine. Western blot analysis confirmed ASNS expression in the RS4;11_ASNS line (**Figure 3A**, left panel). RS4;11_ASNS cells exhibited decreased sensitivity to L-asparaginase (ASNase) treatment compared to the parental line, but increasing concentrations of ASX-173 effectively restored sensitivity to ASNase (**Supplemental figure S3A** and **Figure 3A**, right panel). These findings support the hypothesis that ASX-173 is a specific ASNS inhibitor.

**Figure 3.**
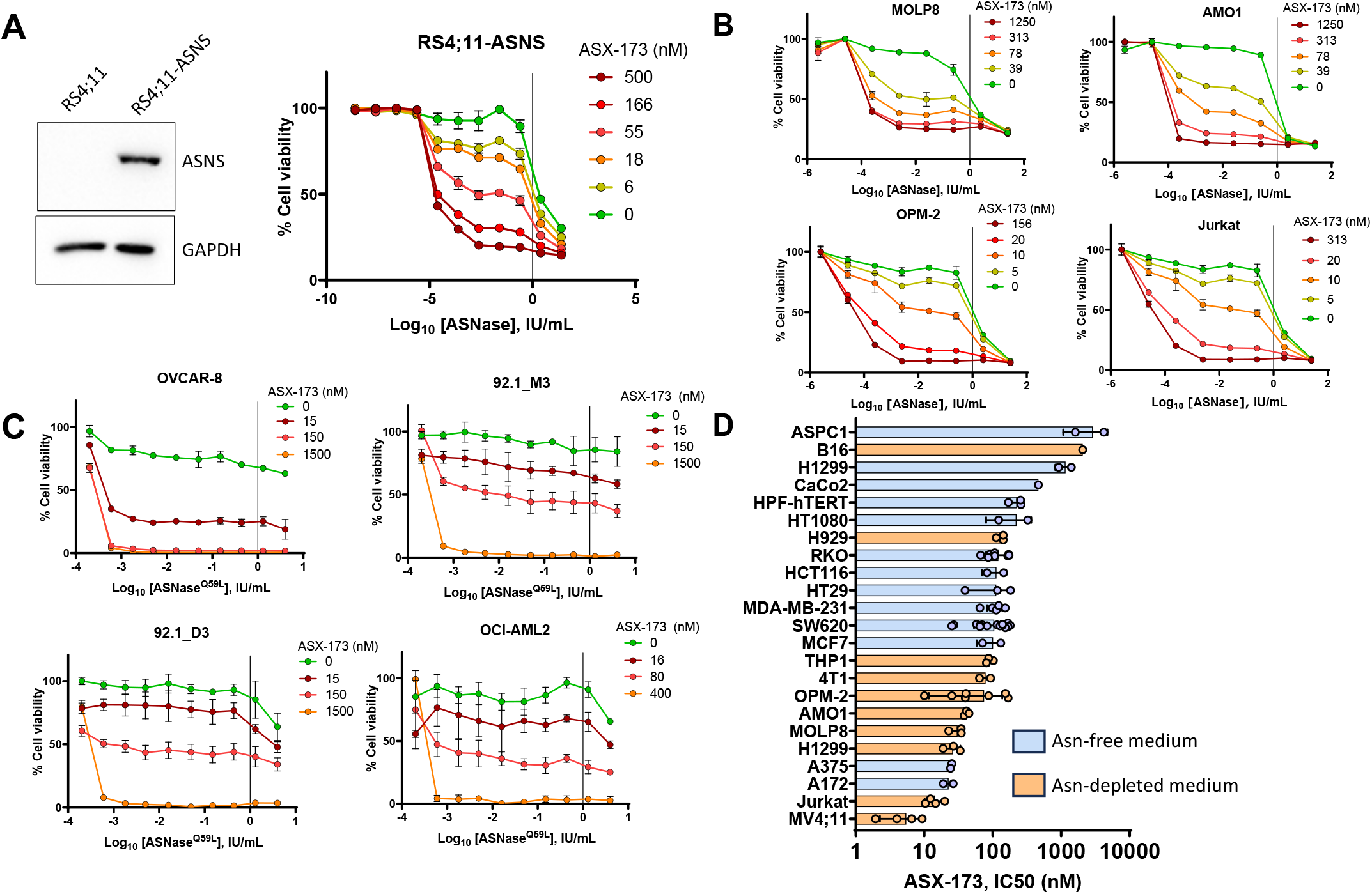
ASX-173 enhances the therapeutic activity of ASNase across multiple cancer types. (**A**) Western blot analysis of ASNS expression in ASNS-negative and ASNS-positive RS4;11 cells. (**B**) ASNS-positive RS4;11 cells were seeded in 96-well plates and treated with a range of ASNase concentrations in combination with the indicated concentrations of ASX-173 for 72 hours. Cell viability was assessed using the Resazurin assay. (**C**) Resazurin assays were performed on two myeloma cell lines (OPM-2, MOLP-8), one plasmacytoma cell line (AMO-1), and one T-cell leukemia cell line (Jurkat) under the same conditions as in panel (**B**). (**D**) Two uveal melanoma cell lines (92.1_M3 and 92.1_D3), one ovarian cancer cell line (OVCAR-8), and one acute myeloid leukemia cell line (OCI-AML2) were treated with a glutaminase-deficient ASNase variant (ASNase_Q59L_) and ASX-173 for 72 hours, as described in panel (**B**). All cell lines, except OCI-AML2, were seeded and incubated overnight before treatment. Cell viability was measured using the CellTiter-Blue assay, with fluorescence excitation at 544 nm and emission at 590 nm. (**E**) IC_50_ values of ASX-173 were determined across various cancer cell lines from different tissue origins under asparagine-deprived conditions (either asparagine-free medium or medium containing 0.025 IU/mL ASNase). Cell viability was assessed using the Resazurin assay for suspension cells and MTT assays for adherent cell lines.

We extended those findings to additional ASNS-positive cancer cell lines, including multiple myeloma (OPM-2, MOLP-8), plasmacytoma (AMO-1), leukemia (Jurkat, H929, MV4;11), and sarcoma (HT1080). In all cases, ASX-173 enhanced the anticancer activity of ASNase (**Figure 3B** and **Supplemental figures S3B**). To distinguish the contributions of the asparaginase and glutaminase activities of ASNase to the observed effects, we next tested ASX-173 in combination with ASNase wild-type (ASNase^WT^) or a glutaminase-deficient ASNase variant (ASNase^Q59L^)^13^ against ASNS-positive cell lines representing ovarian cancer (OVCAR-8), uveal melanoma (92.1_D3, 92.1_M3), and acute myeloid leukemia (OCI-AML2). Although all lines exhibited resistance to ASNase^Q59L^ alone, addition of ASX-173 increased sensitivity in a dose-dependent manner (**Figure 3C and Supplemental figure S3C**), confirming that ASX-173 specifically enhances the anticancer activity associated with ASNase’s asparagine-depleting activity.

Finally, we evaluated the broad-spectrum anticancer activity of ASX-173 against a diverse panel of cancer cell lines under asparagine-deprived conditions, using either asparagine-free medium or low-dose ASNase (0.025 IU/mL). The majority of tested lines (18 of 23) exhibited high sensitivity to the treatments, with ASX-173 IC50 values ranging from 10 to 100 nM (**Figure 3D**), suggesting broad therapeutic potential for ASX-173 in combination with asparagine deprivation.

### ASX-173 induces cell cycle arrest and activates signaling pathways under asparagine-depleted conditions

To probe the cellular mechanisms underlying ASX-173 anticancer activity, we investigated its effects on cell cycle progression in the MV4;11 leukemia cell line. Whereas asparagine-deficient media or ASX-173 treatment alone, even at concentrations up to 500 nM, did not significantly alter cell cycle distribution, combining ASX-173 with asparagine deprivation led to pronounced changes in cell cycle dynamics. Flow cytometry analysis revealed a marked increase in the sub-G1 population accompanied by decreased G2/M and S phase populations, indicating enhanced cell cycle arrest in G1/G0 and apoptotic cell death (**Figure 4A**).

**Figure 4.**
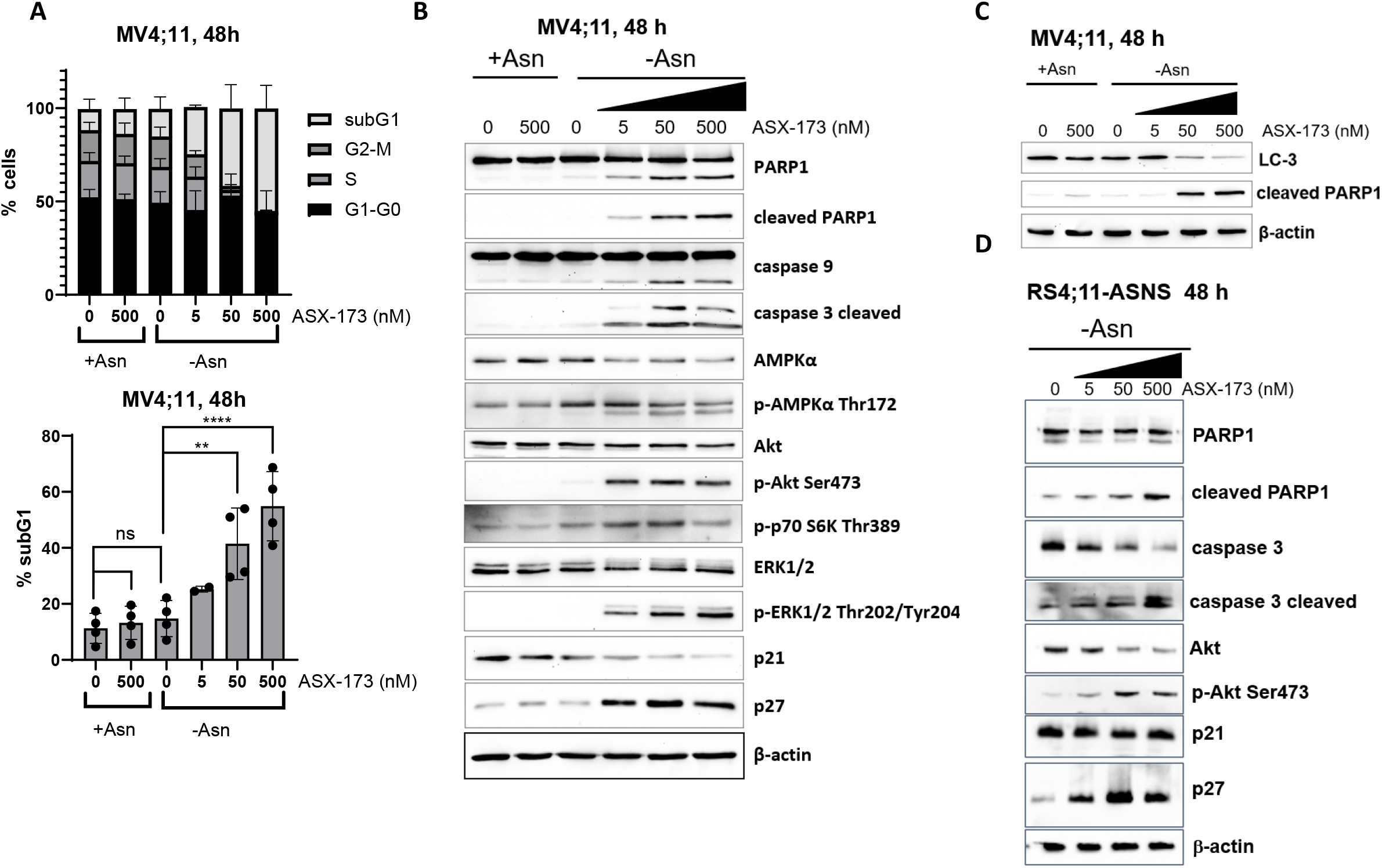
ASX-173 triggers cell cycle arrest and modulates signaling pathways under asparagine deprivation. (**A**) Upper panel: Cell cycle analysis of MV4;11 cells treated with the indicated concentrations of ASX-173 for 48 hours in either asparagine-free or asparagine-supplemented RPMI-1640 medium. Lower panel: Percentage of sub-G1 population from the corresponding cell cycle analysis in the upper panel. Statistical significance was assessed using the ANOVA test with Holm-Šídák’s multiple comparisons test. ** indicates p<0.01, **** indicates p<0.0001 (**B-D**) Western blot analysis of the indicated protein expression levels in cell lysates of MV4;11 cells (**B, D**) and RS4;11-ASNS cells (**C**) treated with indicated concentrations of ASX-173 for 48 hours in either asparagine-free or asparagine-supplemented RPMI-1640 medium. β-actin was used as a loading control.

Molecular analysis of cell cycle and survival pathways provided further insight into the mechanisms underlying the observed anticancer activity. Under asparagine-deprived conditions, ASX-173 treatment triggered a dose-dependent increase in apoptotic markers, including cleaved caspase-3 and cleaved PARP (**Figures 4B-C**). ASX-173 combined with asparagine deprivation also induced distinct changes in the cyclin-dependent kinase inhibitors (CKIs) p27 and p21, which help to maintain cells in non-proliferative states. The treatment upregulated p27, which is a positive modulator of quiescence (reversible G0), and downregulated p21, which is a positive modulator of p53-dependent cell cycle arrest and senescence (irreversible G0). The combination treatment also elicited adaptations in cellular signaling networks. We observed increased expression and activation of pro-survival pathways, indicated by enhanced phosphorylation of AKT, p70S6K, and ERK1/2. Notably, the energy sensor AMPKα was downregulated by the treatment, suggesting a shift toward energy conservation through downregulation of biosynthetic pathways. Consistent with the fact that asparagine depletion is known to activate autophagy^14-17^ we observed downregulation of the autophagic marker LC3 with ASX-173 treatment, suggesting that ASX-173 increases autophagic flux under asparagine-deprived conditions (**Figure 4C**).

Similar signaling responses were observed in RS4;11_ASNS cells (**Figure 4D**), indicating that these adaptations represent a conserved response to combined ASNS inhibition and asparagine depletion.

### ASX-173 and ASNase combination treatment disrupts nucleotide biosynthesis

To delineate the metabolic consequences of combined ASNS inhibition and asparagine depletion, we conducted targeted metabolomic analysis of OCI-AML2 cells with a focus on central energy metabolism pathways and nucleotide metabolism. We compared four treatment conditions: vehicle control, ASNase (0.01 IU/mL), ASX-173 (40 nM), or their combination. The analysis quantified 188 distinct metabolites (see metabolomics raw data), with principal component analysis (PCA) confirming high data quality through tight grouping of biological replicates (**Figure 5A**).

**Figure 5.**
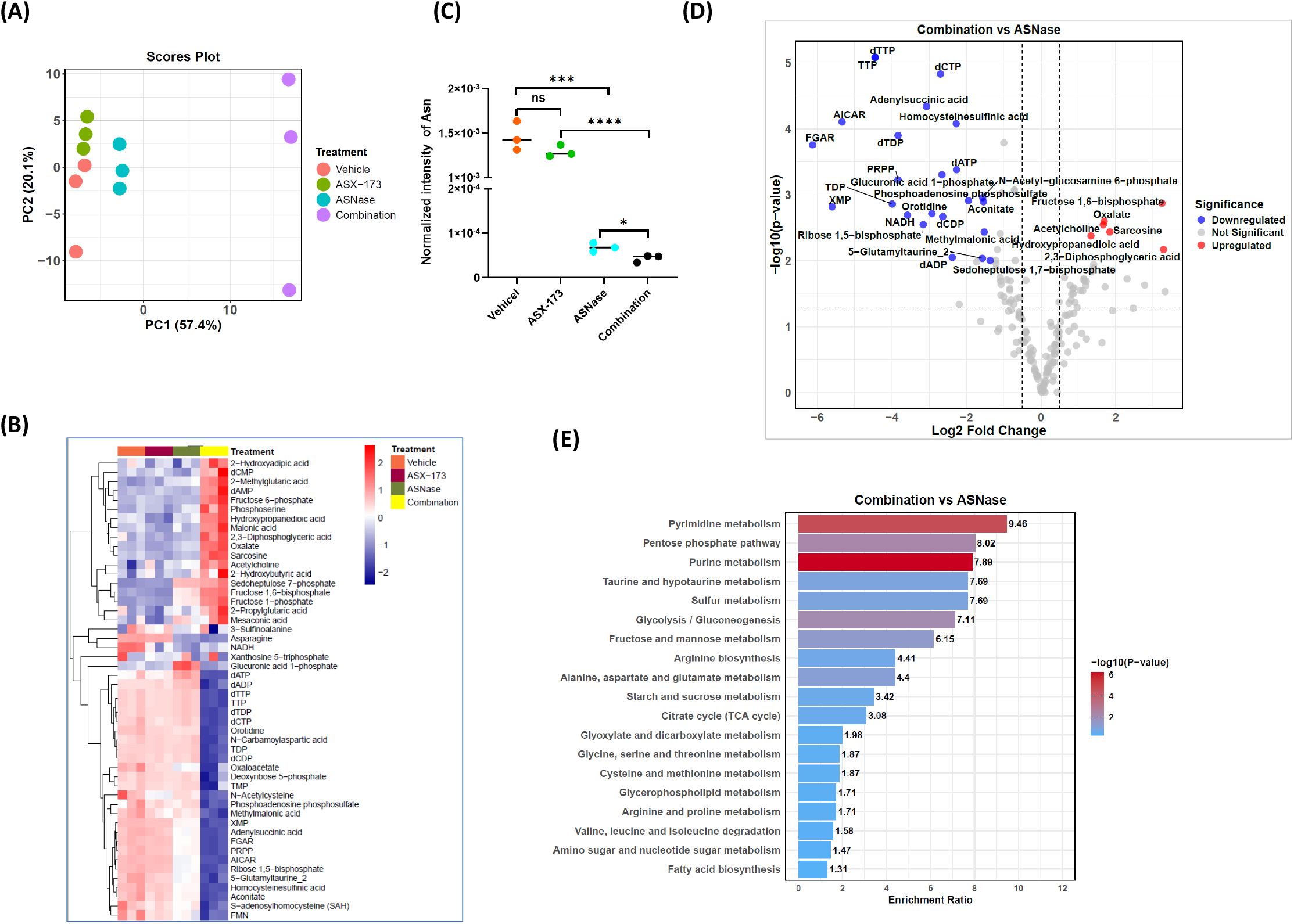
ASX-173 in combination with ASNase disrupts nucleotide biosynthesis. (**A**) Principal Component Analysis (PCA) score plot illustrating the metabolic profiles of OCI-AML2 leukemia cells treated with 0.025 IU/mL ASNase (Spectrila), 80 nM ASX-173, or their combination for 24 hours. Each point represents an individual sample, color-coded by experimental group. The vehicle-treated group serves as a negative control. (**B**) Hierarchical clustering heatmap of metabolite abundance in the samples from (**A**). Rows represent individual samples, while columns correspond to metabolites. Clustering patterns reveal distinct metabolic signatures across treatment groups. (**C**) Relative asparagine levels in each treatment group (n=3) from (**A**), highlighting the impact of ASX-173 and ASNase on asparagine depletion. Statistical significance was assessed by unpaired two-tailed Student’s t-test. *p* < 0.05 (*), *p* < 0.001 (***), *p* < 0.0001 (****), ns = not significant. (**D**) Volcano plot showing metabolite changes between ASNase and Combination groups in (**A**). Each point represents a metabolite; significantly upregulated (Fold change >= 2.5, *p* < 0.01) and downregulated metabolites are highlighted in red and blue, respectively. Non-significant metabolites are shown in gray. (**E**) Bar plot of significantly enriched metabolic pathways based on differentially abundant metabolites between ASNase and Combination groups in (**A**). The top three pathways—pyrimidine metabolism, pentose phosphate pathway, and purine metabolism—indicate that nucleotide biosynthesis is a major affected process in the combination group.

Notably, neither ASX-173 nor low-dose ASNase alone markedly altered metabolite profiles, which clustered closely with those of vehicle-treated cells (**Figures 5A–B**). However, the combination induced significant metabolic reprogramming (**Figure 5B**, top 50 metabolites shown) and slightly greater depletion of intracellular asparagine compared to ASNase treatment alone (**Figure 5C**). Differential analysis revealed no significantly altered metabolites between the ASX-173 and vehicle groups (**Supplemental Figure S5A**). In the ASNase vs. vehicle comparison, six metabolites were significantly downregulated and three were significantly upregulated (fold change ≥ 2.5, p < 0.01, **Supplemental Figure S5B**), indicating minimal metabolic disruption by low-dose ASNase. In contrast, combination treatment resulted in significant downregulation of 36 metabolites and upregulation of 15 compared to vehicle (**Supplemental Figure S5C**). Comparison of the combination treatment to ASNase alone identified 37 significantly altered metabolites, including 25 decreased and 12 increased metabolites (**Figure 5D**). Notably, 17 out of 25 decreased metabolites are directly involved nucleotide metabolism.

Pathway enrichment analysis using the KEGG database provided further evidence that the combination treatment predominantly downregulated nucleotide biosynthesis pathways; the most significantly downregulated pathways (p < 0.001) included pyrimidine metabolism, the pentose phosphate pathway (which produces precursors for nucleotide synthesis), and purine metabolism (**Figure 5E**). These findings suggest that a key mechanism of sensitivity to dual targeting of extracellular asparagine depletion and intracellular ASNS inhibition centers on inhibition of nucleotide biosynthesis.

### ASX-173 potentiates ASNase anticancer efficacy *in vivo*

The therapeutic potential of targeting asparagine metabolism has been limited by rapid development of resistance, with ASNase achieving clinical success only against pediatric acute lymphoblastic leukemia (ALL) despite broad *in vitro* activity. To evaluate whether ASX-173 could overcome that limitation, we tested the combination of ASX-173 and ASNase against the OCI-AML2 leukemia model, which exhibits sensitivity to ASNase *in vitro* but resistance *in vivo*^18^.

Daily treatment with the combination of ASNase (5,000 IU/kg, i.p.) and ASX-173 (50 mg/kg, p.o.) for 14 days effectively suppressed leukemia progression, whereas monotherapy with either agent showed no significant effect (**Figures 6A-B**). The combination therapy achieved a 7-day growth delay over the monotherapies, representing approximately 3-4 leukemia doubling times (**Figure 6B**). The enhanced efficacy of the combination translated to improved overall survival, with all combination-treated mice living 7-10 days longer than those in other treatment groups (**Figure 6C**). Although the combination therapy induced mild toxicity, evidenced by body weight loss (<15%), treated mice fully recovered after treatment cessation (**Figure 6D**).

**Figure 6.**
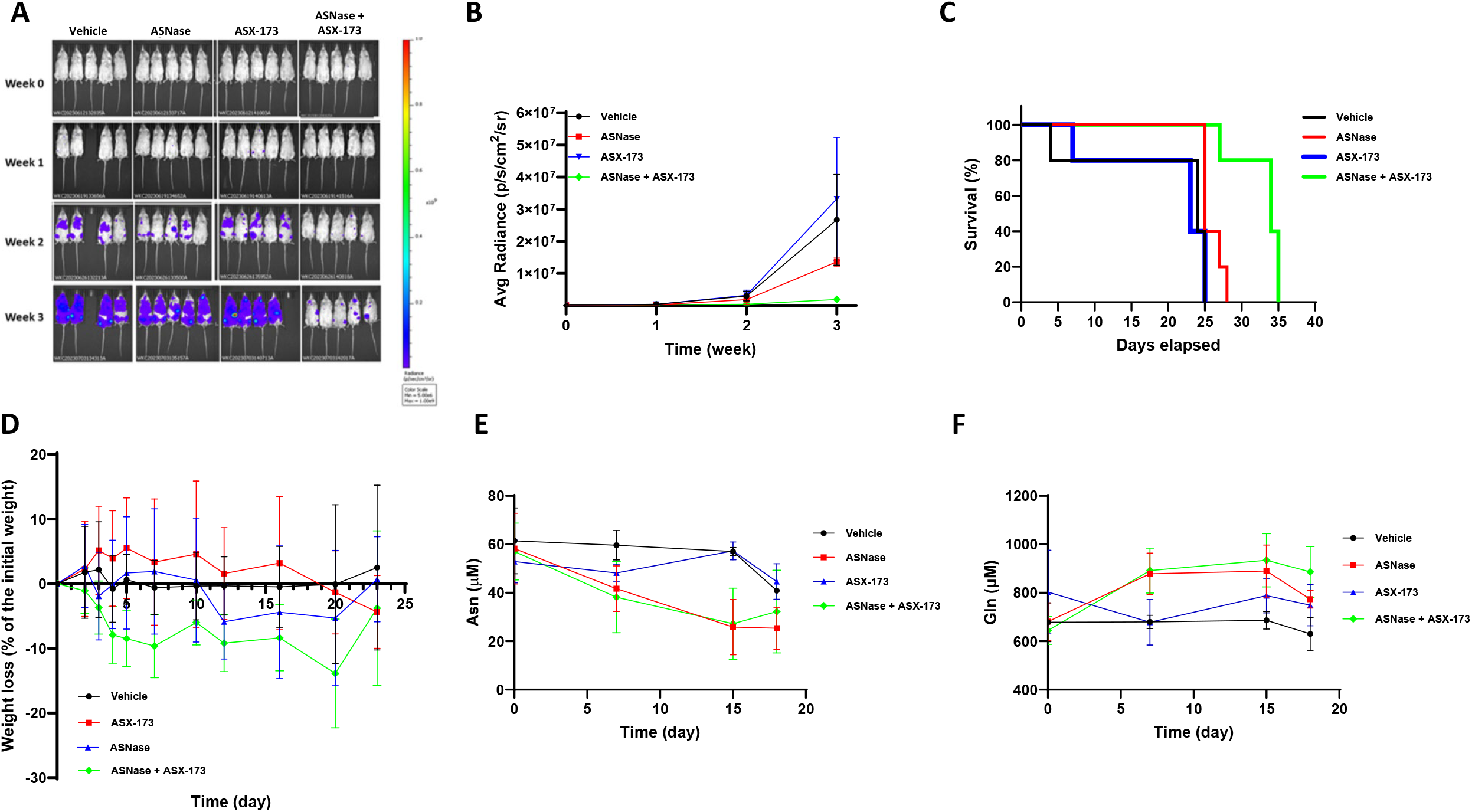
ASX-173 enhances ASNase anticancer efficacy in vivo. NSG mice (n=5 per group) xenografted with luciferase-expressing OCI-AML2 leukemia cells were treated daily for two weeks with PBS (vehicle control), 2,500 U/kg ASNase (Spectrila), 50 mg/kg ASX-173, or their combination. ASNase was administered via intraperitoneal injection, while ASX-173 was given orally. (A) Leukemia burden was monitored using bioluminescence imaging at the indicated time points. Week 0 represents the day before the first treatment. (B) Quantitation of average bioluminescence signal for each treatment group from (A). (C) Kaplan-Meier survival analysis of mice from (A). (D) Average daily body weight change for each treatment group. (E, F) Pharmacodynamic analysis of ASNase in whole blood from mice in (A), measuring asparagine (E) and glutamine (F) concentrations by LC-MS. Mean and standard deviation (SD) are shown.

Pharmacodynamic analysis revealed insight into the mechanistic basis for the therapeutic synergy. Whereas ASX-173 monotherapy did not affect systemic asparagine levels compared to vehicle control, both ASNase alone and the combination treatment significantly decreased plasma asparagine from approximately 60 µM baseline to 25 µM after two weeks (**Figure 6E**). Plasma glutamine levels with ASNase or combination treatment were slightly elevated above baseline (**Figure 6F**), consistent with previous studies and likely due to upregulation of GLS. Notably, aspartate and glutamate levels remained unchanged across all treatment groups (**Supplemental Figure S4**).

### High expression of ASNS is associated with decreased survival of patients with solid tumors

To evaluate the potential clinical utility of ASX-173 beyond hematologic malignancies, we analyzed ASNS expression across various cancer types using TCGA data. ASNS protein expression varied among tumor types (**Figure 7A**), and several cancers, including COAD, EMC, HNSC, LIHC, LUAD, RCC, and UCEC, exhibited significantly higher ASNS protein levels compared to corresponding normal tissues (**Figure 7B**). Furthermore, elevated *ASNS* mRNA expression was associated with poorer patient survival in HNSC, KIRC, and LIHC (**Figures 7C–E**). In KIRC, high ASNS protein levels were also linked to worse survival, supporting the hypothesis that ASNS expression may be a prognostic biomarker across a range of cancer types (**Figures 7F–H**).

**Figure 7.**
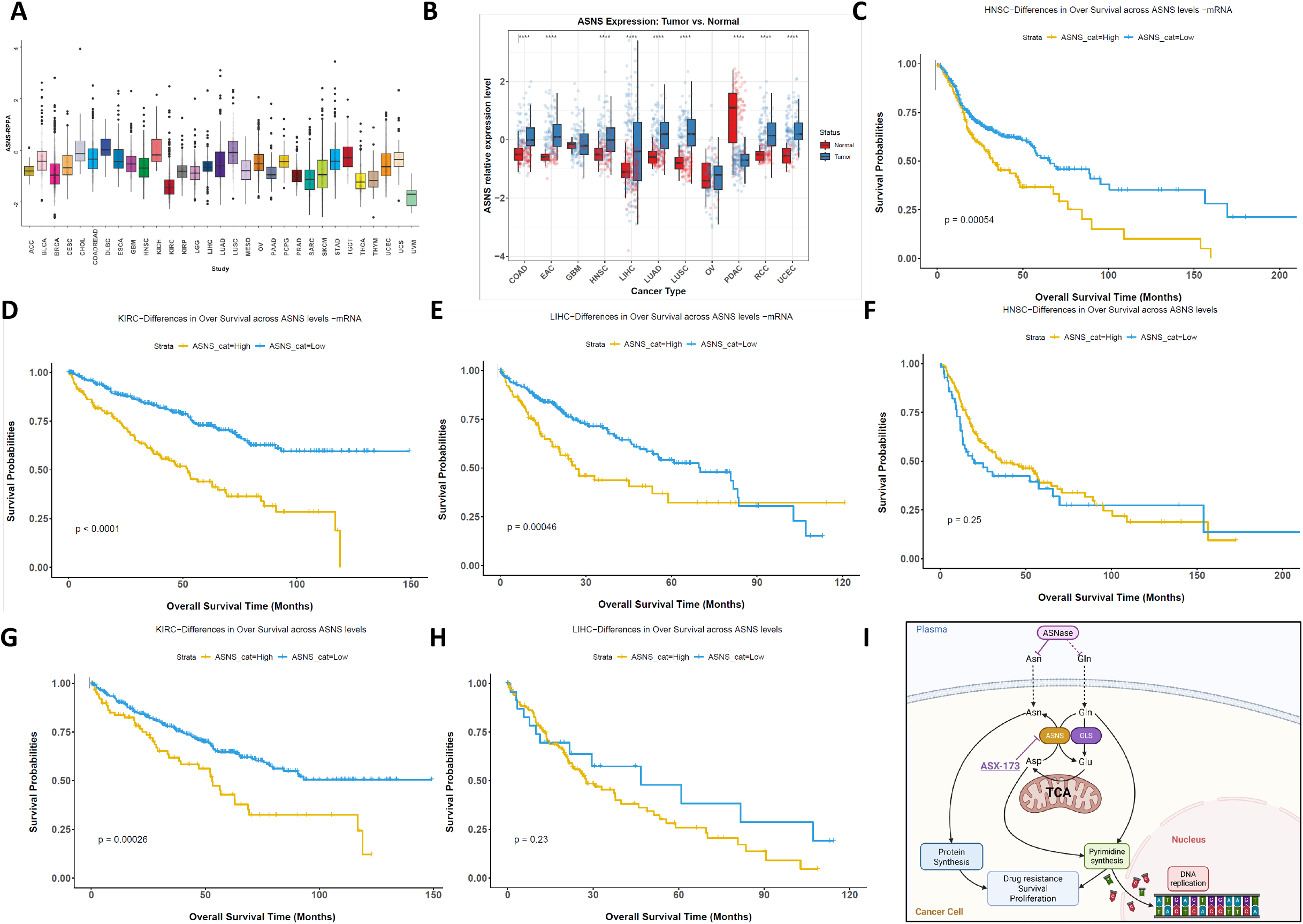
Analysis of ASNS expression in TCGA dataset across various solid tumor types. **(A**) ASNS protein expression across various solid tumor types using TCGA RPPA data. (**B**) ASNS protein expression across indicated tumor and normal tissues in TCGA. Statistical significance was assessed using the Wilcoxon rank-sum test. **** indicates p < 0.0001. (**C**–**E**) Kaplan-Meier survival analysis based on ASNS mRNA expression in selected TCGA cancer types. (**F-H**) Kaplan-Meier survival analysis based on ASNS protein expression in selected TCGA cancer types using RPPA data. Patients were divided into high and low expression groups based on the mean ASNS mRNA level or protein level. Survival differences were assessed using the log-rank test, and *p*-values were shown. HNSC (**C, F**), KIRC (**D, G**) and LIHC (**E, H**). (**I**) Model of ASNase and ASX-173 synergy. Details provided in the main text.

## DISCUSSION

Consistent with previous observations of synthetic lethality between Wnt pathway inhibition and asparagine depletion^19,20^, our high-throughput screen for inhibitors of EMT led to the discovery of effective compounds whose primary targets were ASNS. This finding represents a significant advance for the field, since long-running efforts to develop potent ASNS inhibitors for cancer therapy have fallen short. First-generation ASNS inhibitors, including glutamine analogs^21^, β-aspartyl-adenylate intermediates^22^, and a platinum complex^23^, demonstrated anticancer activity at nanomolar concentrations *in vitro*. However, their *in vivo* efficacy was limited by instability, poor cell permeability, and insufficient specificity that led to dose-limiting toxicities.

From our multidisciplinary approach combining high-throughput screening, kinetic characterization, and cell-based assays, ASX-173 emerged with the greatest specificity and potency for ASNS inhibition. ASX-173 significantly enhanced ASNase anticancer activity, with the combination inducing cell cycle arrest, promoting apoptosis, and inhibiting nucleotide synthesis. Importantly, our preclinical studies demonstrated that the combination was effective *in vivo*; ASX-173 sensitized highly malignant, drug-resistant OCI-AML2 xenografts to ASNase (**Figure 6**).

Our findings provide a rationale for revisiting asparagine metabolism as a target for treatment of a broad range of cancer types. First, ASNS is upregulated across the landscape of cancer (**Figure 7A**). Second, high ASNS expression in the context of cancer is unfavorable for survival in multiple cancer types (**Figures 7B-C**). Third, we recently reported that ASNase treatment rapidly and extensively upregulates ASNS expression in the lung, small intestine, large intestine, liver, and kidney, suggesting that broad ASNS upregulation following ASNase therapy could potentially explain cancer resistance to ASNase treatment^24^. Fourth, we recently found that ASNS also mediates resistance to glutaminase inhibition with CB-839 (Chan WK, et al. under revision), suggesting rational combination of ASX-173 and CB-839. Furthermore, there is a rich literature and long-running history of cell lines reported to be sensitive to ASNase *in vitro* but not *in vivo*—a phenomenon that can be explained by the lack of ASNS inhibition *in vivo*. That is, *in vitro*, the presence of ASNase creates a sink condition that constantly depletes asparagine, effectively inhibiting ASNS. ASX-173 now enables the effective inhibition of ASNS in vivo, which was the missing link to achieving broad ASNase efficacy *in vivo*. Overall, our findings warrant expanded testing of ASX-173 in combination with ASNase and/or glutaminase inhibitors against a broad spectrum of cancer types.

Asparagine was reported to be essential for endothelial cell growth during angiogenesis, a critical process in tumor blood vessel formation^25^. Recent studies have also revealed a strong link between asparagine bioavailability and cancer metastasis^26^. Proteomic analyses indicated that asparagine enrichment is a conserved feature of proteins driving epithelial-to-mesenchymal transition (EMT), suggesting that asparagine may influence metastasis by regulating EMT. Given the close link between angiogenesis and metastasis, this relationship may explain why elevated ASNS expression is consistently associated with poor outcomes across various cancer types, including worse prognoses in colorectal cancer and increased aggressiveness in lung cancer and glioblastoma. These findings further support a rationale for testing the combination of ASX-173 and ASNase against metastatic and aggressive cancers.

A recent study identified bisabosqual A (Bis A) as a novel ASNS inhibitor that covalently modifies the K556 site of ASNS^27^. However, its anticancer activity has not been validated *in vivo*, and concerns remain about potential irreversible toxicities even after treatment cessation due to covalent ASNS modification. In contrast, ASX-173 monotherapy exhibits a favorable safety profile, with no significant changes in metabolomic profiles *in vitro* or weight loss *in vivo*. In addition, the reported *in vitro* synergy of Bis A and ASNase was weaker than that of the ASX-173 and ASNase. Notably, the study did not include rescue experiments with asparagine to confirm the specificity of Bis A.

Our findings suggest additional combination strategies for future investigation. The activation of ERK1/2 and AKT signaling in response to ASX-173 treatment points to potential synergies with MAPK and PI3K/AKT pathway inhibitors. The impact on nucleotide metabolism also suggests combinations with DNA-damaging agents or other compounds targeting nucleotide synthesis pathways.

Because synergistic efficacy is usually accompanied by synergistic toxicity, a notable limitation of the current study is the absence of comprehensive toxicity assessments. Additional animal toxicity studies and a Phase I clinical study will be planned to assess the safety profile of the combination of ASX-173 and ASNase.

Our analysis of TCGA data underscores the broader therapeutic potential of ASX-173 beyond hematologic malignancies by revealing elevated ASNS expression across several cancer types, suggesting that ASNS upregulation may represent a common metabolic adaptation in diverse tumor settings. These findings support the rationale for targeting ASNS in a wider range of malignancies and highlight the clinical relevance of ASX-173 as a promising therapeutic candidate, particularly in KIRC, where high ASNS expression is associated with poor clinical outcomes.

In conclusion, ASNS plays a critical role in both protein and pyrimidine synthesis, not only by catalyzing the production of asparagine but also by acting as a glutaminase to generate glutamate, which feeds into downstream pyrimidine synthesis. This dual function is essential for cancer cell survival, drug resistance to asparaginase (ASNase), metastasis, and proliferation (**Figure 7I**). ASX-173 represents a significant advance in targeting asparagine metabolism for cancer therapy. In combination with ASNase, the resulting bicompartmental blockade exhibits a broad spectrum of anticancer activity, a clear mechanism of action, and *in vivo* efficacy. Future studies to optimize pharmacokinetics, test rational combinations, and examine additional cancer models will help to realize the full therapeutic potential of targeting asparagine metabolism for the treatment of cancer.

## MATERIALS AND METHODS

### Compounds and Materials

ASX-173 was synthesized by Asinex, with synthesis details provided in the Supplementary Methods. Commercial *E. coli* ASNase was sourced from Wacker Biotech (Spectrila) or Lens-Farm, while recombinant ASNase^Q59L^ was produced as previously described. Antibodies for AMPKα, p-AMPKα, AKT, p-AKT, ASNS, CASPASE 9, ERK1/2, p-ERK1/2, p21, p27, and β-ACTIN were purchased from Cell Signaling and are listed in **Supplemental Table S1**. RPMI-1640, DMEM, and NEAA were obtained from Invitrogen or Paneko.

### Determination of asparagine synthetase activity

The activity of recombinant wild-type human asparagine synthetase in the presence of the tested compound was assessed by measuring the rate of inorganic pyrophosphate (PPi) production using a continuous assay with Pyrophosphate Reagent (Sigma). This assay couples PPi production to NADH consumption, enabling real-time monitoring of enzyme activity. The details are described in the supplemental materials.

### Cell Culture and Assays

All cell lines were maintained in RPMI-1640, EMEM, IMDM or DMEM supplemented with 5% fetal bovine serum and 2 mM glutamine, as described previously.

TCF/LEF Reporter Assay: SW620 and HCT116 cells were transduced with a TCF/LEF Reporter lentiviral firefly luciferase construct (SABiosciences) and a control Renilla luciferase construct. Cells (10,000/well) were seeded in 96-well black, clear-bottom plates (Corning 3904) and incubated overnight. Cells were treated with 0.02% DMSO or test compounds for 24 hours. Firefly and Renilla luciferase signals were measured using the Dual-Glo® Luciferase Assay System (Promega), with firefly luciferase activity normalized to Renilla. IC_50_ values were determined using a four-parameter variable slope model in GraphPad Prism 8.0.

MTT Assay: Cells were seeded in 96-well black, clear-bottom plates. Adherent cells were allowed to attach overnight before treatment with ASX-173, ASNase, or their combination, while suspension cells were treated immediately. After 72 hours, MTT reagent (Sigma, M5655) was added (0.5 mg/mL) and incubated for 4 hours. Plates were centrifuged (550 x g, 12 min, 4°C), supernatants removed, and 100 μL of DMSO added per well. Plates were shaken for 1 hour at room temperature, and absorbance was measured at 570 nm. Blank wells (without MTT) were used for background subtraction. Control wells were set to 100%, and IC_50_ values were calculated using GraphPad Prism 8.0.

Resazurin Assay: Cells were seeded and treated as in the MTT assay. After 72 hours, resazurin reagent (Sigma, R7017) was added (40 μM) and incubated for 4-6 hours. Fluorescence was measured (Ex: 560 nm, Em: 590 nm). IC_50_ values were determined as in the MTT assay.

CellTiter-Blue Cell Viability Assay was performed as previously described.

### Generation of Firefly Luciferase-Overexpressing OCI-AML2 Cell Line

Lentiviral particles were produced by transiently transfecting HEK293T cells with a firefly luciferase expression plasmid along with the packaging plasmids RSV-Rev, pMDLg/pRRE, and CMV-VSVG, using jetPRIME Transfection Reagent (#101000046, Polyplus) according to the manufacturer’s instructions. Viral supernatants were collected 48 hours post-transfection and used to infect OCI-AML2 cells in the presence of 8 μg/mL polybrene.

To generate the firefly luciferase-overexpressing OCI-AML2 cell line, infected cells were cultured for 48 hours and subsequently sorted for green fluorescent protein (GFP) expression via flow cytometry. The GFP-positive cell population was expanded in culture for two weeks. 1 × 10^6^ GFP-positive cells were then engrafted into NSG mice (NOD.Cg-PRKDC(scid) IL2RG(tm1Wjl); The Jackson Laboratory stock #005557) via tail vein injection.

One week post-engraftment, leukemia burden was assessed using bioluminescence imaging (BLI) with an IVIS Imaging System (PerkinElmer). Blood was collected from the mouse exhibiting the highest BLI signal, and OCI-AML2 cells were isolated by flow cytometry. The GFP-positive cell population was further expanded through another round of *in vivo* selection to establish a stable firefly luciferase-expressing OCI-AML2 cell line.

### Real-Time PCR

Cells were seeded in 6 cm Petri dishes and incubated overnight. The following day, cells were treated with 0.02% DMSO or ASX-173 for 24 hours. RNA was extracted using Trizol reagent (ThermoFisher), quantified by NanoDrop, and reverse-transcribed using Superscript II Reverse Transcriptase (Invitrogen). qRT-PCR was performed with SYBR Green PCR Mix (Evrogen) on a Bio-Rad Thermal Cycler. Relative gene expression was analyzed using the ΔΔCt method, with GAPDH as the reference gene. Primer sequences are listed in **Supplemental Table S2**.

### Western Blotting

MV4;11, RS4;11, and RS4;11-ASNS cells were seeded in 10 cm Petri dishes in RPMI-1640 or asparagine-free RPMI-1640 and treated with 0.02% DMSO or various ASX-173 concentrations for 48 hours. Cells were centrifuged (1000 x g, 5 min, 4°C), washed with ice-cold PBS, and lysed in RIPA buffer (containing 2 mM PMSF and a protease inhibitor cocktail) for 30 min on ice. Protein concentrations were determined using the Bradford assay. Lysates (40 μg protein) were separated by SDS-PAGE, transferred to 0.2 μm nitrocellulose membranes (Bio-Rad), and blocked with 5% skim milk in TBST. Membranes were incubated overnight at 4°C with primary antibodies (Supplementary Table S2), followed by HRP-conjugated secondary antibodies for 1 hour at room temperature. Proteins were visualized using Clarity Western ECL Substrate (Bio-Rad) and imaged with the iBright FL1500 Imaging System (Invitrogen). Representative immunoblots and images are presented.

### Flow Cytometry

MV4;11 cells were treated as in Western blotting. After 48 hours, cells were centrifuged (1000 x g, 5 min, 4°C), washed with PBS, and stained with propidium iodide (50 μg/mL) in buffer containing 100 μg/mL RNase A, 0.1% sodium citrate, and 0.3% NP-40. Samples were incubated in the dark for 30 minutes and analyzed using a Cytoflex flow cytometer (Beckman Coulter). At least 10,000 events per sample were recorded and analyzed with CytExpert software (Beckman Coulter).

### Metabolomics

OCI-AML2 cells (1.2 × 10^6^) were treated with vehicle, 0.025 IU/mL ASNase (Spectrila), 80 μM ASX-173, or their combination in RPMI-1640 for 24 hours. Cells were centrifuged (250 x g, 1 min, 4°C), and metabolites were extracted using methanol:water (80:20, v/v) + 0.1% ammonium hydroxide. After processing, samples were analyzed by ion chromatography-mass spectrometry (IC-MS) and liquid chromatography-mass spectrometry (LC-MS). The details are provided in the Supplementary Methods. Data were acquired using a Thermo Orbitrap Fusion Tribrid Mass Spectrometer and analyzed with MetaboAnalyst.

### Mouse Leukemia Model

Mouse studies were performed in a pathogen-free vivarium at The University of Texas MD Anderson Cancer Center under an approved Institutional Animal Care and Use Committee (IACUC) study protocol (ACUF #00001658-RN00). NSG mice (n=5 per group) were injected with 0.5 × 10^6^ luciferase-engineered OCI-AML2 cells via the tail vein. After two weeks, leukemia burden was monitored by bioluminescence imaging (IVIS, PerkinElmer). Mice were randomized into four groups: PBS (control), 2,500 IU/kg Spectrila, 50 mg/kg ASX-173, or their combination. Treatments were administered for two weeks, and leukemia burden was assessed weekly via imaging as described previously.

### DepMap analysis

CRISPR Chronos dependency data was obtained from the DepMap Public 24Q4 dataset (https://depmap.org/portal/). Information on culture conditions was downloaded from the DepMap data portal (https://depmap.org/portal/data_page), and cell lines were classified into NEAA+ and NEAA-groups. The list of genes essential for each media type was generated using Depmap two-class comparison analysis, which calculates groups means for sensitivity scores to knockout in the CRISPR Chronos dataset. All the downloaded data are provided in the supplemental file of **ASX-173_Depmap_Analysis.xlsx**.

### Data Analysis

Statistical analyses were conducted using GraphPad Prism 8. Group differences were evaluated using one-way ANOVA. Details on sample size (n), statistical methods, and error bar definitions are provided in the figure legends. Metabolomic data were analyzed using MetaboAnalyst 6.0 ^28^. Raw data and additional statistics are provided in supplemental file **ASX-173_Metabolomics.xlsx**. All experiments were repeated at least twice unless otherwise specified. Representative immunoblots and images are included. ASNS TCGA data were download from cBioPortal (www.cbioportal.org) or The Human Protein Atlas (www.proteinatlas.org). Raw data are provided in supplemental file **ASX-173_TCGA_ASNS.xlsx**.

## Author contributions

V.K., W.K.C., D.G., R.K. and P.L.L. designed the project; R.K and D.G. performed chemical design, V.K., W.K.C., N.J., K.A., P.N., D.A. performed cell culture work; W.K.C., L.T., S.A.M. and B.Q.T. performed metabolomics experiments; W.K.C., L.T., I.M., S.A.M. and B.Q.T. performed metabolomics data analysis; S.V.K performed TCGA data analysis. A.S. and M.K. generated firefly luciferase-overexpressing OCI-AML2 cell line; W.K.C. performed *in vivo* animal experiments; V.K., W.K.C., L.T., J.N.W., R.K., and P.L.L. wrote the manuscript.

## Acknowledgments

This work was supported by Asinex Corporation. This work was supported in part by NCI grant numbers CA143883 (J.N.W), CA083639 (J.N.W.), CA235510 (J.N.W.), and CA016672 (J.N.W. and P.L.L.); Cancer Prevention and Research Institute of Texas grant number RP130397 (J.N.W. and P.L.L.); The Metabolomics Core Facility is also supported by NIH grants S10OD012304-01 and P30CA016672.

## Declaration of interests

The authors declare the following financial interest(s): Roman Kombarov and Dmitry Genis are employees of Asinex Corporation. ASX-173 was synthesized by Asinex Corporation. Victor Tatarskiy was a part-time employee of Asinex Corporation at the time this work was carried out. The other authors declare no conflict of interest.

## Supplemental Materials

Supplemental Methods, Supplemental Figures S1-S5; Supplemental Tables S1 and S2, Raw data files: ASX-173_Depmap_analysis.xlsx, ASX-173_Metabolomics.xlsx and ASX-173_TCGA_ASNs.xlsx

## Notes

### Competing Interest Statement

Roman Kombarov and Dmitry Genis are employees of Asinex Corporation. ASX173 was synthesized by Asinex Corporation. Victor Tatarskiy was a part-time employee of Asinex Corporation at the time this work was carried out. The other authors declare no conflict of interest.

