## Supplemental methods, Supplemental Tables and Supplemental Figures for "The ASNS inhibitor ASX-173 potentiates L-asparaginase anticancer activity"

### Supplementary Methods

#### I. Synthesis of ASX-173

NMR spectra were registered on «MERCURY plus 400 MHz» spectrometer (Varian). Chemical shift values are given in ppm relative to tetramethylsilane (TMS), with the residual solvent proton resonance as internal standard. The test samples were dissolved in DMSO-$d_6$ or $CDCl_3$. To the extent NMR peak forms and multiplicities are specified, they are stated as they appear in the spectra, possible higher order effects have not been considered.

Analytical LCMS spectra were recorded on two instruments at the following conditions:

Instrument **A**: Surveyor MSQ (Thermo Fisher Scientific)
Column:Phenomenex Onyx Monoliythic C18 25 x 4.6 mm Part No: CHO-7645.
LC eluent A: 0.1% of formic acid in water, LC eluent B: 0.1% of formic acid in acetonitrile
Gradient: a linear gradient of B from 5% to 100%for 5 min.
Flow rate: 1.5 ml/min.
Injection volume: 2µL
Detection UV: PDA -photodiode array detector in 200-800 nm range
Detection Mass: APCI (+ or – ions) - atmospheric pressure chemical ionization.

Instrument **B**: Agilent Infinity 1260, Agilent 6120 (Agilent Technologies Inc.)
Column:Kinetex 5uM EVO C18 100A 50 x 4.6 mm column
LC eluent A: 0.1% of formic acid in water, LC eluent B: 0.1% of formic acid in acetonitrile
Gradient: a linear gradient of B from 5% to 100%for 5 min.
Flow rate: 1.3 ml/min.
Injection volume: 2 µL
Detection UV: diode array (Agilent DAD1260; G4212B), at 275 nm
Detection Mass: APCI (+ or – ions) - atmospheric pressure chemical ionization.

Preparative HPLC purifications were performed in reverse phase mode on HPLC system equipped with a Jasco PU-2086 Plus preparative pump, Rheodyne Model 7125 Injector, Thermo Hypersil-Keystone BETASIL™ PREP C18 HPLC column P/N 70110-259270 Dim. 250×21.2 mm, particle size 21 µm, and Agilent Multi-Wavelength Detector (G1365B).The instrument was operated by Agilent ChemStation B.04.03 software in Microsoft Windows 7. LC eluent A: 0.1 % HCOOH in water, LC eluent B: 0.1 % HCOOH in acetonitrile, flow rate at 20 mL/min, UV detection at 210 nm, injection volume: 500 µL.

Column chromatography was done by using Kieselgel 60 (Merck) 60–200 mesh as the stationary phase.

All reagents and solvents were purchased from commercial sources and used without further purification. All yields reported in this publication refer to isolated ones of compounds and their purity was determined by 1H NMR and LCMS.

The stereochemistry displayed in the products is relative and not absolute.

**Scheme 1. Synthesis of Intermediate 10**

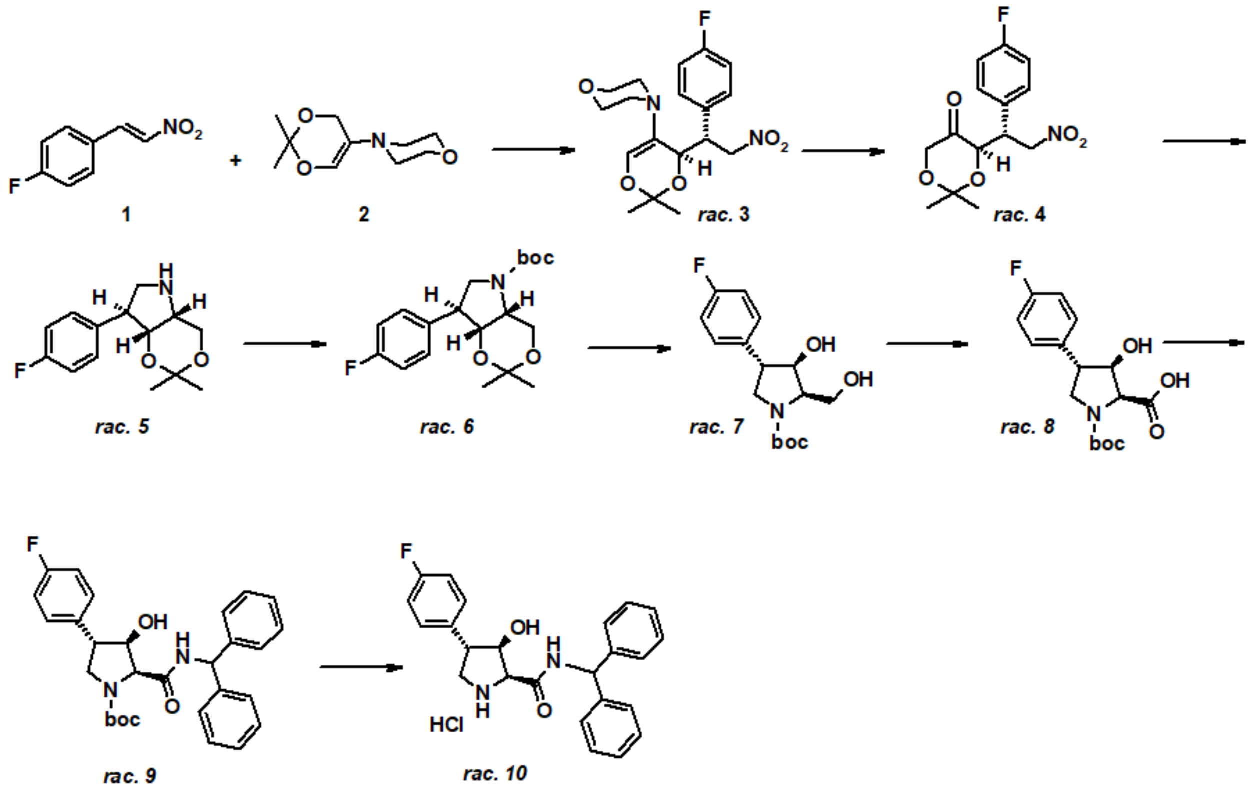


**rac-4-[(4R*)-4-[(1R*)-1-(4-fluorophenyl)-2-nitroethyl]-2,2-dimethyl-2,4-dihydro-1,3-dioxin-5-yl]morpholine (3).**

A stirred solution of1-fluoro-4-(2-nitrovinyl)benzene (77.79 g, 0.465 mol) and 2,2-dimethyl-5-morpholino-4H-1,3-dioxin(77.28 g, 0.388 mol) in acetonitrile (720 mL) was heated at 40-45ºC for 12 hours. The solvent was removed *in vacuo*, and the residue was treated with methanol (150 mL). The formed white crystals were collected by filtration, washed with cold methanol and air-dried to afford the title product (102.8 g, 72%).

$^1$H NMR (400 MHz, $CDCl_3$) δ 7.36 (dd, $J$ = 8.6, 5.6 Hz, 2H), 6.94 (t, $J$ = 8.7 Hz, 2H), 4.90 (dd, $J$ = 13.3, 8.3 Hz, 1H), 4.71 – 4.59 (m, 2H), 3.99 (td, $J$ = 7.7, 3.0 Hz, 1H), 3.79 – 3.61 (m, 4H), 2.74 (ddd, $J$ = 11.5, 6.2, 3.3 Hz, 2H), 2.21 (ddd, $J$ = 11.4, 6.2, 3.1 Hz, 2H), 1.46 (2x s, 6H).

**rac-(4R*)-4-[(1R*)-1-(4-fluorophenyl)-2-nitroethyl]-2,2-dimethyl-1,3-dioxan-5-one (4).**

To a solution of **3** (101.8 g, 0.278 mol) in acetonitrile (1.1 L) was added at a solution of oxalic acid dihydrate (35.02 g; 0.278 mol, 1.0 eq.) in water (450 mL) and the reaction mixture was stirred at 40-42ºC for 6 hours. A second portion of oxalic acid dihydrate (3.16 g; 25 mmol, 0.1 eq.) was added and the reaction mixture was stirred at 40-42ºC for 4 hours, then allowed to stand overnight at rt. Upon reaction completion, the mixture was treated with $NaHCO_3$ (4.6 g), NaCl (30 g) and ethyl acetate (100 mL). The mixture was stirred at rt for 15 min. The organic phase was separated and concentrated *in vacuo*. The residue was extracted with dichloromethane (200 mL), and the extract was dried over $Na_2SO_4$ and concentrated i*n vacuo*. The residue was purified by flash-chromatography (hexane/ethyl acetate 4:1) to afford the title product as a light-yellow oil (77.2 g, 93%).

$^1$H NMR (400 MHz, $CDCl_3$) δ 7.31 (dd, $J$ = 8.6, 5.5 Hz, 2H), 6.98 (t, $J$ = 8.6 Hz, 2H), 4.87 (dd, $J$ = 12.8, 8.4 Hz, 1H), 4.65 (dd, $J$ = 12.8, 7.1 Hz, 1H), 4.57 (d, $J$ = 3.7 Hz, 1H), 4.13 (ddd, $J$ = 8.4, 7.1, 3.8 Hz, 1H), 3.89 (s, 2H), 1.46 (2x s, 6H).

**rac-(4aS*,7R*,7aS*)-7-(4-fluorophenyl)-2,2-dimethylhexahydro-2H-[1,3]dioxino[5,4-b]-**

**pyrrole (5).**

A solution of **4** (77g; 0.259 mol) in methanol (1.2 L) was hydrogenated in 2 L Parr steel autoclave over Raney nickel (150 g) under 10 bar at rt. Upon reaction completion, the reaction mixture was filtered through Celite®. The filtrate was concentrated *in vacuo*, and the residue was dissolved in dichloromethane (150 mL) and dried over $Na_2SO_4$. The solvent was removed *in vacuo* to afford the title product as a brown oil (56.2 g, 86%).

$^1$H NMR (400 MHz, $CDCl_3$) δ 7.22 (dd, $J$ = 8.6, 5.4 Hz, 2H), 7.01 (t, $J$ = 8.6 Hz, 2H), 4.19 (d, $J$ = 3.1 Hz, 1H), 4.13 (dd, $J$ = 12.7, 3.8 Hz, 1H), 3.95 (dd, $J$ = 12.6, 2.8 Hz, 1H), 3.73 (dd, $J$ = 11.7, 8.1 Hz, 1H), 3.34 (dd, $J$ = 8.1, 5.0 Hz, 1H), 3.12 – 3.03 (m, 2H), 1.42 (2x s, 6H).

***rac*-tert-butyl (4aS*,7R*,7aS*)-7-(4-fluorophenyl)-2,2-dimethyl-hexahydro-2H-[1,3]dioxino[5,4-b]pyrrole-5-carboxylate (6).**

To a solution of **5** (56.2 g, 0.224 mol) in dichloromethane (400 mL) was added a solution of $K_2CO_3$ (42.04 g, 0.304 mol) in water (250 mL). To the resulting mixture, a solution of di-*tert*-butyl dicarbonate (48.75 g, 0.224 mol) in dichloromethane (150 mL) was added dropwise at stirring over 20 min at rt. The mixture was stirred for 12 hours, the organic layer was separated and dried over $Na_2SO_4$. The solvent was removed *in vacuo*, and the residue was purified by flash-chromatography on silica gel, eluting with hexane-ethyl acetate 4:1 to afford the title product as a light-brown oil which was used further without additional purification (55.0 g, 70%).

***rac*-tert-butyl (2*R**,3*R**,4*S**)-4-(4-fluorophenyl)-3-hydroxy-2-(hydroxymethyl)-pyrrolidine-1-carboxylate (7).**

A mixture of **6** (55 g; 0.157 mol) in acetic acid (207 mL) and 89 mL of water was stirred at 45ºC for 5 hours. Then the mixture was diluted with water (800 mL) and the product was extracted with ethyl acetate (3 x 250 mL). The combined organic extracts were washed with 10% $Na_2CO_3$ and dried over $Na_2SO_4$. The solvent was removed *in vacuo* and the residue was treated with hexane (100 mL). The solid material was collected by filtration and dried at 60ºC for 2 hours to provide the title product as a white solid (42.18 g, 87%).

$^1$H NMR (400 MHz, $CDCl_3$) δ 7.22 (dd, $J$ = 8.4, 5.3 Hz, 2H), 7.02 (t, $J$ = 8.6 Hz, 2H), 4.41 (t, $J$ = 7.7 Hz, 1H), 4.17 – 4.04 (m, 2H), 4.00 – 3.86 (m, 2H), 3.49 (dd, $J$ = 10.8, 9.2 Hz, 1H), 3.40 (q, $J$ = 8.5 Hz, 1H), 2.71 (br s, 2H), 1.14 (s, 9H).

***rac*-(2*S**,3*R**,4*S**)-1-[(tert-butoxy)carbonyl]-4-(4-fluorophenyl)-3-hydroxypyrrolidine-2-carboxylic acid (8).**

A solution of **7** (42.1 g; 0.135 mol) and TEMPO (1.56 g; 10 mmol) in acetonitrile (680 mL) and phosphate buffer (pH 6.7; 590 mL) was heated to 40-45ºC. To this heated solution a solution of of 80% sodium chlorite (36.7 g; ~0.32 mol) in water (160 mL) and 3% sodium hypochlorite "bleach" (6.3 mL) in water (63 mL) were simultaneously added from two funnels over 15 min. The resulting reaction mixture was stirred at 40-45ºC for 10 hours. Upon cooling to rt, $Na_2SO_3$ (33.6 g) and NaCl (122 g) were added, and pH was adjusted to 4 by adding 4M HCl. The organic layer was separated, and the aqueous layer was extracted with THF (3 x 100 mL). The combined organic extracts were concentrated *in vacuo* and the residue was dissolved in the mixture of dichloromethane/THF (4:1) (300 mL). The residual water layer was discarded, and the organic layer was dried over $Na_2SO_4$ and concentrated *in vacuo*. The solid residue was treated with MTBE (80 mL) and collected by filtration, washed with MTBE and dried at 50ºC to provide the title product as a white solid (26.7 g, 61%).

$^{1}$H NMR (400 MHz, DMSO) δ 7.28 – 7.23 (m, 2H), 7.11 (td, *J* = 8.5, 5.1 Hz, 2H), 4.47 (t, *J* = 8.4 Hz, 1H), 4.32 (2x d, *J* = 7.9 Hz, 1H), 3.87 (m, 1H), 3.43 – 3.26 (m, 2H), 1.15 (s, 9H).

***rac*-tert-butyl (2S*,3R*,4S*)-2-[(diphenylmethyl)carbamoyl]-4-(4-fluorophenyl)-3-hydroxypyrrolidine-1-carboxylate (9).**

A mixture of **8** (1.00 g; 3.1 mmol), TBTU (1.14 g; 3.6 mmol) and DIPEA (1.14 mL; 6.4 mmol) in acetonitrile (20 mL) was stirred at rt for 20 minutes. Then benzhydrylamine (0.62 g; 3.4 mmol) was added, and the reaction mixture was stirred at rt for 8 hours. The solvent was removed *in vacuo*, the residue was partitioned between dichloromethane and 10% $K_2CO_3$, and the obtained emulsion was filtered. The organic layer of the filtrate was separated, washed with 10% citric acid, water and dried over $Na_2SO_4$. The solvent was removed *in vacuo*, and the residue was treated with hexane/MTBE 1:1 (20 mL). The formed solid was collected by filtration and air-dried to provide the title product as a white solid (1.32 g, 87%).

***rac*-(2*S**,3*R**,4*S**)-N-(diphenylmethyl)-4-(4-fluorophenyl)-3-hydroxypyrrolidine-2-carboxamide hydrochloride (10)**.

A mixture of **9** (1.32 g, 2.7 mmol) in 18% HCl in dioxane (50 mL) was stirred at rt for 30 min. The volatiles were removed *in vacuo*, and the residue was triturated with diethyl ether. The formed precipitate was collected by filtration, washed with diethyl ether, and air-dried to provide the title product as a white amorphous solid (1.13 g, 98%).

$^{1}$H NMR (400 MHz, DMSO) δ 8.44 (d, *J* = 8.4 Hz, 1H), 7.37 – 7.18 (m, 12H), 7.09 (t, *J* = 8.9 Hz, 2H), 6.11 (d, *J* = 8.4 Hz, 1H), 5.04 (br s, 1H), 4.25 (q, *J* = 5.2 Hz, 1H), 3.72 (d, *J* = 6.1 Hz, 1H), 3.43 (dd, *J* = 10.4, 7.3 Hz, 1H), 3.12 (td, *J* = 7.0, 4.7 Hz, 1H), 2.86 (dd, *J* = 10.4, 6.6 Hz, 1H).

**Scheme 2**

**(9H-fluoren-9-yl)methyl N-[(2S)-1-[(2S,3R,4S)-2-[(diphenylmethyl)carbamoyl]-4-(4-fluorophenyl)-3-hydroxypyrrolidin-1-yl]-1-oxobutan-2-yl]carbamate (11a); 9H-fluoren-9-ylmethyl (9H-fluoren-9-yl)methyl N-[(2S)-1-[(2R,3S,4R)-2-[(diphenylmethyl)carbamoyl]-4-(4-fluorophenyl)-3-hydroxypyrrolidin-1-yl]-1-oxobutan-2-yl]carbamate (11b).**

To a stirred solution of **10** (480 mg, 1 eq.), (2S)-2-(9H-fluoren-9-ylmethoxycarbonylamino)-butanoic acid (366 mg, 1 eq.) and DIPEA (465 µL, 2.5 eq.) in dry DMF (9 mL), TBTU (435 mg, 1.2 eq.) was added in one portion. The resulting mixture was stirred at rt for 25 min. The reaction mixture was subjected to HPLC separationon using Thermo Hypersil-Keystone BETASIL™ PREP C18 HPLC column P/N 70110-259270, eluting with gradient of 0.1% formic acid in acetonitrile/ 0.1% formic acid in water 60%-70% at 20 mL/min for 30 min, collecting fractions by UV at 210 nm, to provide **11a** (retention time: 24.9 min) and **11b** (retention time: 26.6 min). Upon fraction evaporation the title products were obtained as white foams (**11a** 154 mg, **11b** 390 mg).

**Scheme 3**

**(2R,3S,4R)-1-[(2S)-2-aminobutanoyl]-N-(diphenylmethyl)-4-(4-fluorophenyl)-3-hydroxypyrrolidine-2-carboxamide (ASX-173)**

To a stirred solution of **11b** (390 mg) in dry DMF (4 mL), DBU (400 µL) was added. The resulting mixture was stirred at rt for 30 min. The solvents were removed in vacuo and the crude product was purified by column chromatography on silica gel, eluting with gradient of ethyl acetate / methanol 1:0, 1:0.1; 1:0.2 to provide the title products as a white solid (225 mg, 85%).

mp 180-181 °C;

rotation value [α]20D (10g/L, DCM) = 4.8;

APCI-MS m/z calcd. for C28H30FN3O3 475, found 476 $[M+H]^+$;
1H NMR (DMSO-d6, 400 MHz) 8.81 (d, J = 8.2 Hz, 1/2H), 8.47 (d, J = 8.2 Hz, 1/2H), 7.38–7.16 (m, 30 12H), 7.11 (m, 2H), 6.13 (t, J = 8.0 Hz, 1H), 5.55 (br. s, 1/2H), 5.28 (br. s, 1/2H), 4.83 (d, J = 7.7 Hz, 1/2H), 4.61 (d, J = 7.8 Hz, 1/2H), 4.42 (m, 1H), 4.13 (t, J = 9.0 Hz, 1/2H), 3.81 (dd, J = 11.6, 8.9 35 Hz, 1/2H), 3.53 (m, 1H), 3.39 (m, 1/2H) 3.28 (t, J = 10.9 Hz, 1/2H), 3.14 (dd, J = 7.8, 5.3 Hz, 1H), 1.62 – 1.46 (m, 1/2H), 1.29 (m, 1H), 1.08 (m, 1/2H), 0.85 (t, J = 7.4 Hz, 3/2H), 0.50 (t, J = 7.4 Hz, 3/2H).

#### II. Determination of ASNS enzymatic activity

The reagent couples the production of inorganic pyrophosphate to NADH consumption. The assay mixture contained 5 mM ATP, 10 mM L-aspartate, 100 mM $NH_4Cl$ and 10 mM $MgCl_2$ dissolved in 100 mM EPPS buffer, pH 8, and either 10 nM, 30 nM, 50 nM or 80 nM of the tested compound (1 mL total volume). Reactions were initiated at 25°C by the addition of 8 µL of a solution containing hASNS to a final concentration of 10 nM hASNS. NADH consumption was monitored spectrophotometrically at 340 nm over a period of 40 min. All kinetic assays were performed in duplicates and repeated two times. The curve figures represent the mean values of duplicate experiments. It was assumed that the binding of the nine inhibitors would be competitive with respect to ATP, according to the following kinetic model:

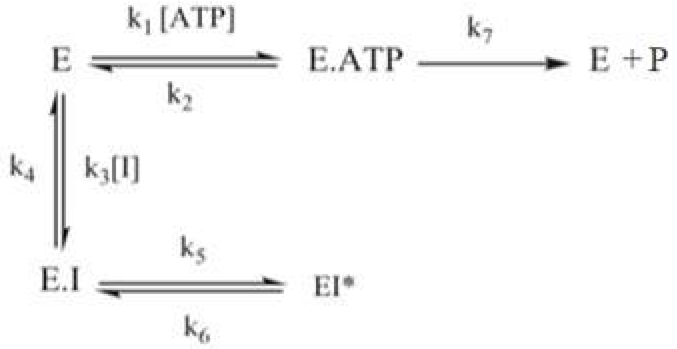


Scheme 1a. The *k*s were estimated by fitting the curve to the following equation:
$[PPi] = v_{ss}t + (v_0 - v_{ss})*(1 - e^{-kt})/k$
where [PPi] is the concentration of inorganic pyrophosphate formed at time $t$, $v_o$ and $v_{ss}$ are the initial and steady-state rates, respectively, and $k$ is the apparent first-order rate constant for isomerization of EI to EI*
$k_5$, $k_6$ and $K_i$ were estimated by fitting the plots of $k$ vs [I] using the following equation:

$$k = k_6 + k_5\left(\frac{I/Ki}{1+\frac{A}{ka}+I/Ki}\right)$$

$K_i$* was calculated as
$K_i^* = K_i k_6/(k_6+k_5)$

#### III. Analysis of Amino Acids and TCA cycle metabolites by LC/IC-MS

Metabolites were extracted using 1 mL ice-cold 0.1% Ammonium hydroxide in 80/20 (v/v) methanol/water. Extracts were centrifuged at 17,000 g for 5 min at 4°C, and supernatants were transferred to clean tubes, followed by evaporation to dryness under nitrogen.

Dried extracts were reconstituted in deionized water, and 5 µL was injected for analysis by ion chromatography (IC)-MS. IC mobile phase A (MPA; weak) was water, and mobile phase B (MPB; strong) was water containing 100 mM KOH. A Thermo Scientific Dionex ICS-6000+ system included a Thermo IonPac AS11 column (4 µm particle size, 250 x 2 mm) with column compartment kept at 30°C. The autosampler tray was chilled to 4°C. The mobile phase flow rate was 360 µL/min, and the gradient elution program was: 0-5 min, 1% MPB; 5-25 min, 1-35% MPB; 25-39 min, 35-99% MPB; 39-49 min, 99% MPB; 49-50, 99-1% MPB. The total run time was 55 min. To assist the desolvation for better sensitivity, methanol was delivered by an external pump and combined with the eluent via a low dead volume mixing tee. Data were acquired using a Thermo Orbitrap IQ-X Tribrid Mass Spectrometer under ESI negative ionization mode at resolution of 240,000.

For amino acids analysis, same samples were injected for analysis by liquid chromatography (LC)-MS. LC mobile phase A (MPA) was 95/5 (v/v) water/acetonitrile containing 20 mM ammonium acetate and 20 mM ammonium hydroxide (pH~9), and mobile phase B (MPB) was acetonitrile. Thermo Vanquish LC system included a Xbridge BEH Amide column (3.5 µm particle size, 100 x 4.6 mm) with column compartment kept at 35°C. The autosampler tray was chilled to 4°C. The mobile phase flow rate was 300 µL/min, and the gradient elution program was: 0-1 min, 85% MPB; 1-16 min, 85-5% MPB; 16-20 min, 5% MPB; 20-21 min, 5-85% MPB. The total run time was 25 min. Data were acquired using a Thermo Orbitrap Exploris 240 Mass Spectrometer under ESI positive/negative ionization (polarity switching) mode at a resolution of 240,000.

Raw data files were imported to Thermo Trace Finder 5.1 and Skyline Daily software for final analysis. The relative abundance of each metabolite was normalized by DNA concentration.

### Supplementary Tables

#### Supplemental Table S1: List of the antibodies used in this study

| Name of target | Clone | Resource / Isotype | Catalog # | Source |
|---|---|---|---|---|
| AMPKα | D63G4 | Rabbit | 5832 | CST |
| Phospho-AMPKα (Thr172) | 40H9 | Rabbit | 2535 | CST |
| Akt (pan) | 11E7 | Rabbit | 4685 | CST |
| Phospho-Akt (Ser473) XP® | D9E | Rabbit | 4060 | CST |
| Phospho-p70 S6 Kinase (Thr389) | D5U1O | Rabbit | 97596 | CST |
| p44/42 MAPK (Erk1/2) | 137F5 | Rabbit | 4695 | CST |
| Phospho-p44/42 MAPK (Erk1/2) (Thr202/Tyr204) XP® | D13.14.4E | Rabbit | 4370 | CST |
| p27 Kip1 XP® | D69C12 | Rabbit | 3686 | CST |
| p21 Waf1/Cip1 | E2R7A | Rabbit | 37543 | CST |
| LC3A/B XP® | D3U4C | Rabbit | 12741 | CST |
| PARP1 | polyclonal | Rabbit | 9542 | CST |
| Cleaved PARP1 (Asp214) XP® | D64E10 | Rabbit | 5625 | CST |
| Caspase-3 | D3R6Y | Rabbit | 14220 | CST |
| Cleaved Caspase-3 (Asp175) | 5A1E | Rabbit | 9664 | CST |
| Caspase-9 | C9 | Mouse | 9508 | CST |
| β-actin | AC-74 | Mouse | A2228 | Sigma-Aldrich |
| Anti-rabbit IgG, HRP-linked | - | Goat | 7074 | CST |
| Anti-mouse IgG, HRP-linked | - | Horse | 7076 | CST |

*CST – Cell Signaling Technology

**Supplementary Table S2: List of the real-time PCR primers used in this study**

| | Primer | Sequence |
|---|---|---|
| 1 | DKK1-F5 | 5'-TTTCCTCAATTTCTCCTCGG-3' |
| 2 | DKK1-R5 | 5'-ATGCGTCACGCTATGTGCT-3' |
| 3 | AXIN2-F6 | 5'-CCCGAGAGCCGGGAAATAAAA-3' |
| 4 | AXIN2-R6 | 5'-TCCAGTTCCTCTCAGCAATCG-3' |
| 5 | CD44-F10 | 5'-CGTGGAATACACCTGCAAAG-3' |
| 6 | CD44-R10 | 5'-CGGACACCATGGACAAGTTT-3' |
| 7 | CD133-F13 | 5'-TTCTTGACCGACTGAGACCCA-3' |
| 8 | CD133-R13 | 5'-TCATGTTCTCCAACGCCTCTT-3' |
| 9 | MYC-F14 | 5'-TGAGGAGACACCGCCCAC-3' |
| 10 | MYC-R14 | 5'-CAACATCGATTTCTTCCTCATCTTC-3' |

### Supplementary Figures

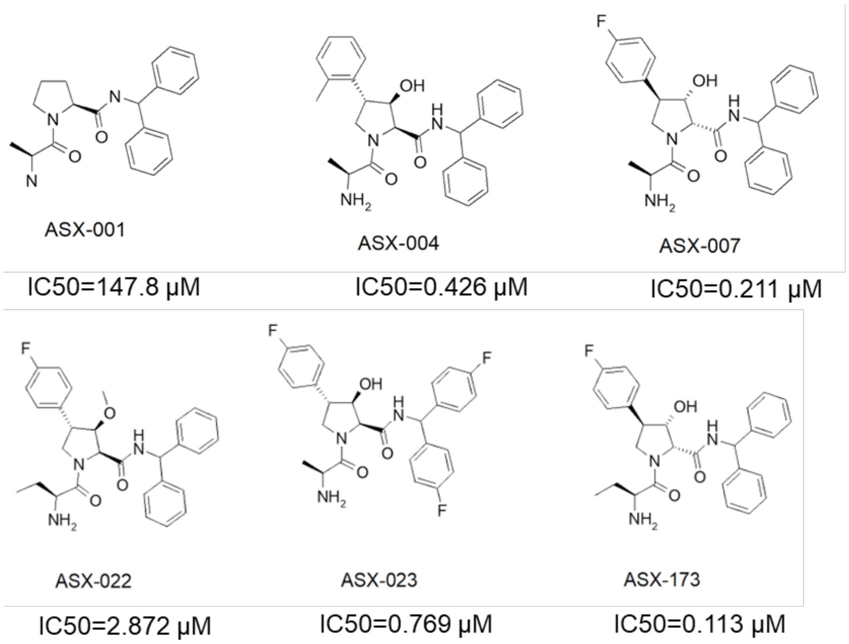


**Supplemental Figure S1. Small molecules identified as positive hits in phenotypic screening.** Chemical structures and their IC50 values were shown.

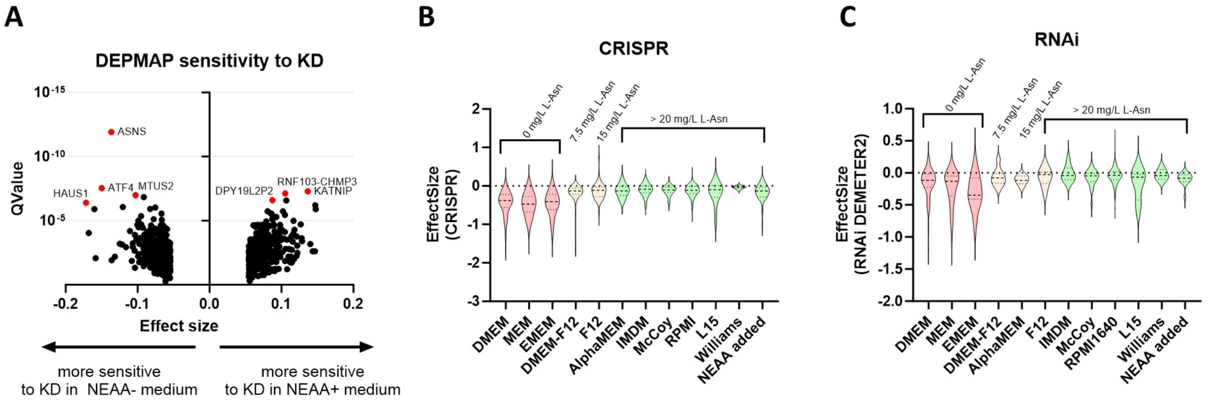


**Supplemental Figure S2.** Analysis of cell line sensitivity to gene knockdown by RNAi in NEAA(**-**) or NEAA(+) medium using DepMap. (A) Volcano plot depicting gene dependency in human cancer cell lines (RNAi knockdown) cultured in NEAA-deficient medium, based on DepMap data. NEAA(+) medium was used as a control. (**B-C**) Sensitivity violin plot of ASNS-deficient cancer cells generated either with CRISPR knockout (**B**) or RNAi knockdown (**C**) to various culture media, as observed in the DepMap database. Asparagine containing medium was used as a control.

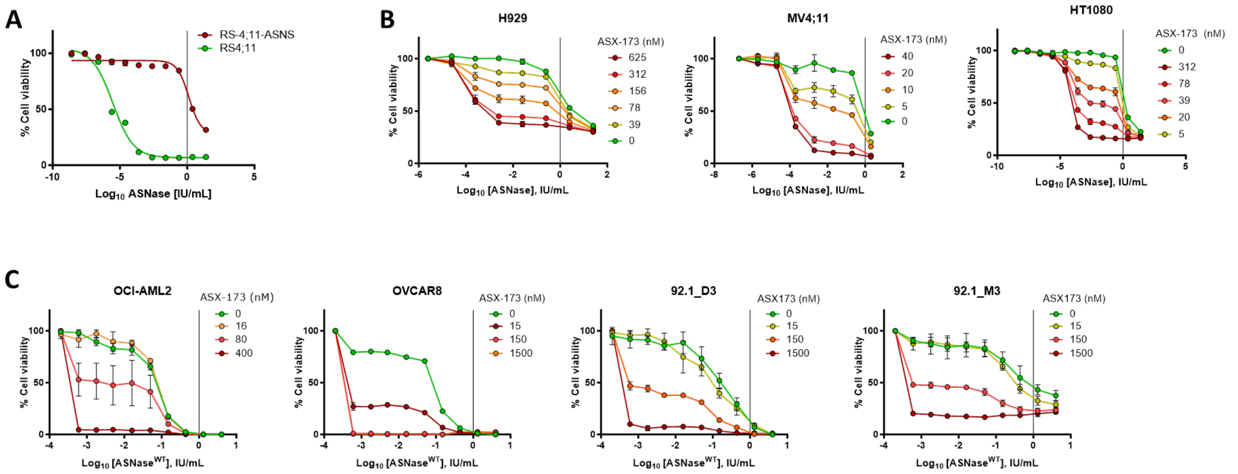


**Supplemental Figure S3**. ASX-173 enhances the therapeutic activity of ASNase across multiple cancer types. (A) RS4;11 and RS4;11-ASNS cells were seeded in 96-well plates and treated with a range of ASNase concentrations in combination with the indicated concentrations of ASX-173 for 72 hours. Cell viability was assessed as in **Figure 3A**. (**B**) Cell viability assays were performed on two leukemia cell lines (H929 and MV4;11) and one sarcoma cell line (HT1080), as described in **Figure 3B**. (**C**) Cell viability assays were performed on two uveal melanoma cell lines (92.1_M3 and 92.1_D3), one ovarian cancer cell line (OVCAR-8), and one acute myeloid leukemia cell line (OCI-AML2), as described as in **Figure 3C**. Cells were treated with ASNase wild-type (ASNase$^{WT}$) and ASX-173 for 72 hours.

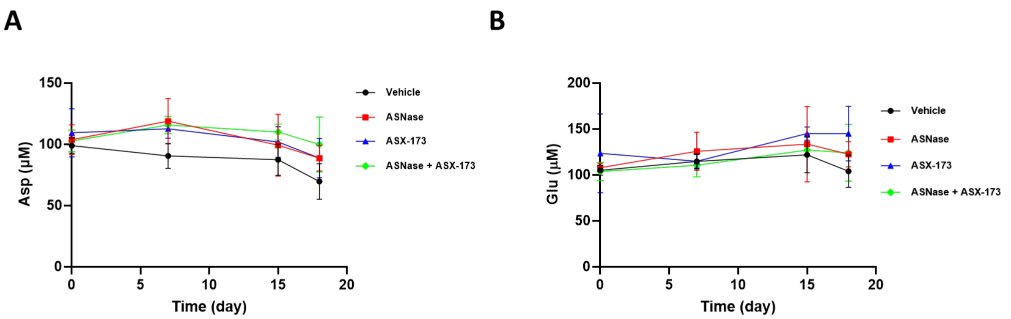


**Supplemental Figure S4.** Pharmacodynamic analysis of ASNase in whole blood from mice. Aspartate (**A**) and glutamate (**B**) concentrations were measured by LC-MS, as described in Figure 6. Mean and standard deviation (SD) are shown.

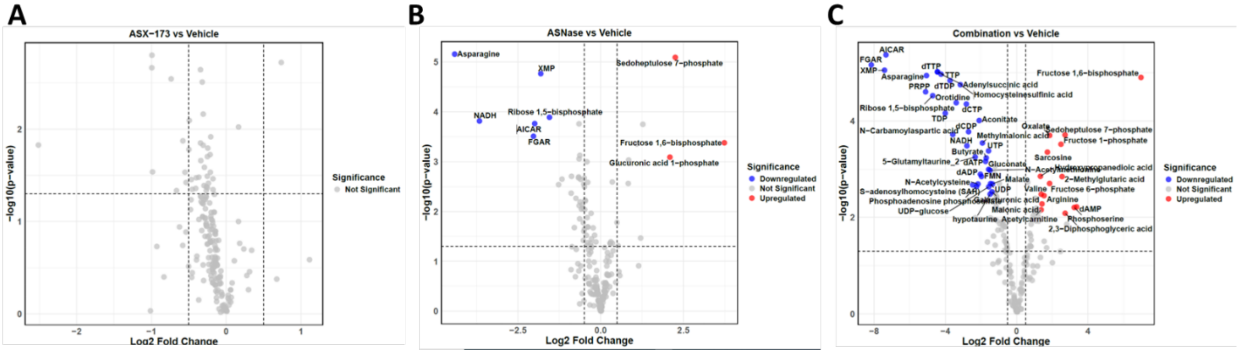


**Supplemental Figure S5.** Volcano plots depicting metabolite changes in ASX-173 vs. Vehicle **(A)**, ASNase vs. Vehicle **(B)**, and Combination vs. Vehicle **(C)** groups. Each point represents an individual metabolite. Significantly upregulated and downregulated metabolites are (fold change ≥ 2.5, $p < 0.01$) highlighted in red and blue, respectively. Metabolites without significant changes are shown in gray.
